# High KIR diversity in Uganda and Botswana children living with HIV

**DOI:** 10.1101/2024.12.03.626612

**Authors:** John Mukisa, Samuel Kyobe, Marion Amujal, Eric Katagirya, Thabo Diphoko, Gaseene Sebetso, Savannah Mwesigwa, Gerald Mboowa, Gaone Retshabile, Lesedi Williams, Busisiwe Mlotshwa, Mogomotsi Matshaba, Daudi Jjingo, David P. Kateete, Moses L. Joloba, Graeme Mardon, Neil Hanchard, Jill A. Hollenbach, the Collaborative African Genomics Network (CAfGEN) of the H3Africa Consortium

**Author notes:** **Corresponding author**: John Mukisa, Department of Immunology and Molecular Biology Makerere University College of Health Sciences P.O.BOX 7072, Kampala, Uganda.

## Abstract

Killer-cell immunoglobulin-like receptors (*KIR*s) are essential components of the innate immune system found on the surfaces of natural killer (NK) cells. The *KIR*s encoding genes are located on chromosome 19q13.4 and are genetically diverse across populations. *KIR*s are associated with various disease states including HIV progression, and are linked to transplantation rejection and reproductive success. However, there is limited knowledge on the diversity of *KIR*s from Uganda and Botswana HIV-infected paediatric cohorts, with high endemic HIV rates. We used next-generation sequencing technologies on 312 (246 Uganda, 66 Botswana) samples to generate *KIR* allele data and employed customised bioinformatics techniques for allelic, allotype and disease association analysis. We show that these sample sets from Botswana and Uganda have different KIRs of different diversities. In Uganda, we observed 147 vs 111 alleles in the Botswana cohort, which had a more than 1 % frequency. We also found significant deviation towards homozygosity for the *KIR3DL2* gene for both rapid (RPs) and long-term non-progressors (LTNPs)in the Ugandan cohort. The frequency of the bw4-80I ligand was also significantly higher among the LTNPs than RPs (8.9 % Vs 2.0%, P-value: 0.032). In the Ugandan cohort, *KIR2DS4*001* (OR: 0.671, 95 % CI: 0.481-0.937, FDR adjusted Pc=0.142) and *KIR2DS4*006* (OR: 2.519, 95 % CI: 1.085-5.851, FDR adjusted Pc=0.142) were not associated with HIV disease progression after adjustment for multiple testing. Our study results provide additional knowledge of the genetic diversity of *KIR*s in African populations and provide evidence that will inform future immunogenetics studies concerning human disease susceptibility, evolution and host immune responses.

## Introduction

Killer-cell immunoglobulin-like receptors (*KIRs*) are crucial modifiers of body immune defense mechanisms found on natural killer cells (NK),^1^, CD4, CD8 and γδ T cells^2–4^. Genes located in the leukocyte-receptor complex region of chromosome 19q13.4 encode the different *KIR*s^1, 5–7^. *KIR*s exhibit high allelic diversity and gene content complexity with two haplotypes (A and B) designated based on the presence or absence of inhibiting and activating genes^8, 9^. *KIR* haplotypes consist of 4-13 genes with high levels of polymorphisms resulting in over 1617 alleles identified so far^1, 10, 11^. This is mainly attributable to several factors. First, there is a high frequency of structural variants due to complex allele and gene content combinations that result from recombination hotspots at the telomeric and centromeric ends^12^. In addition, there are a variable number of insertions and deletions as well as tandem duplications of *KIR* genes^10, 12, 13^. Given this diversity, researchers have the potential to categorize *KIR* gene and allele patterns, discover new alleles across populations, and understand mechanisms of disease susceptibility and survival among individuals.

Previous studies have described the *KIR* gene content and allelic diversity in the Yucpa^14^, Europeans and Asians^15, 16^, Maori and Polynesians^15, 17^, Japanese^18^, European Americans^19, 20^, Malaysians^21^, Han Chinese^22^, Iranians^23^ and Amerindians^24^. Nemat-Gorgani *et al.* have documented high *KIR3DL2* allele frequencies among three African populations^18^. Other studies among the Ga-Adangbe West Africans^25, 26^, Tunisians^27^, Zimbabweans^28^, and Egyptians^29^ have revealed novel alleles at different frequencies between African populations and European populations. These studies also indicate allelic and haplotype diversity of the *KIR* genes between different global and African populations. More immunogenetics-focused research in this region is needed due to the high genetic diversity on the continent^30–34^. To date, most of the studies in Uganda have used non-next-generation sequence techniques which may be limited to the presence or absence of *KIR* genes^35, 36^ due to the complexity of the region, while no studies have been performed in Botswana.

Despite the complexity of the *KIR* region, the advent of new affordable, rapid, and computationally efficient techniques and algorithms for determining the extent and characteristics of the genome at the *KIR* locus has improved our knowledge of immunogenetics^37^. This offers new opportunities for deciphering determinants of complex traits at the genetic level. Specifically, the structural and functional distinctions of various *KIRs* have allowed for a better understanding of their interaction with sequence motifs of Human Leukocyte Antigen (HLA) class I molecules during body defense mechanisms^7^. *KIRs* function by delivering inhibitory and/or activating signals which shape innate immune system responses^1, 7, 19, 38^. This crucial role has been noted in their association with infectious diseases like human immunodeficiency virus (HIV)^33, 39–45^ and malaria^46^, variable clinical presentation of neurological disorders like multiple myeloma and Parkinson’s disease^47^, gastric cancer^48^, preeclampsia^35^, and transplantation disorders^49, 50^. Particularly in HIV, the activated *KIR3DS1* gene in combination with its HLA-C ligand triggers the NK cell to destroy infected cells expressing Major Histocompatibility Class (MHC) class I ligands via degranulation and an increase in cytotoxic T-cell activity^51^, while the inhibitory *KIR3DL1* leads to the dampening of NK cell activity^43, 52^. This mechanism has been postulated to slow disease progression among HIV-infected individuals in studies outside of Africa^40^. However, as demonstrated previously, only a handful of studies focused on HIV disease progression and *KIRs* have been performed among Eastern and southern African populations^53^ known to bear a disproportionately higher burden of HIV^33, 53^. Additionally, children with HIV are a unique vulnerable group with a maturing immune system and varied manifestations of innate and adaptive immune responses^54^. An analysis of the *KIR* loci of African populations offers a foundation for future studies on the evolution of *KIR*, their HLA ligand pairs, and documentation of the disease mechanisms across populations.

Here, we present an evaluation of the *KIR* and HLA class I genotypes and alleles to characterize their diversity in children living with HIV from Botswana and Uganda using the extended Pushing Immunogenetics into the Next Generation (PING) pipeline^37^. We provide one of the first studies to describe *KIR* allele diversity at high resolution and potential *KIR* association with HIV disease phenotypes in high HIV burden settings.

## Materials and Methods

### Study design, population, and ethical considerations

This retrospective cohort study analyzed data from children enrolled in the Collaborative African Genomics Network (CAfGEN) studies as described previously^33, 34, 55^. In brief, the CAfGEN consortium consisted of sites in Uganda, Eswatini, and Botswana at the paediatric centers of excellence in HIV care that collaborated with the University of Botswana, Makerere University, and Baylor College of Medicine, Texas, USA. Clinical and demographic information of the children enrolled through the electronic medical records have previously been described^33, 56^. The Long-term non-progressors (LTNPs) were children who had been asymptomatic for more than 10 years after perinatal HIV-1 infection with a CD4 count above 500 cells/ml or CD4 counts above 25%. Rapid progressors included children with two or more consecutive CD4 below 15%, an AIDS-defining illness (WHO stage III), and antiretroviral therapy initiated within the first 3 years of life after perinatal HIV infection^57, 58^. We performed library preparation and targeted short-read sequencing for *KIR* genes of 349 samples from the *CAfGEN* cohorts recruited in Botswana and Uganda that have previously been described^33, 34, 55^. Meta data for the participants like sex and disease phenotype status were retrieved from the electronic medical records at the HIV care centres of excellence in Botswana and Uganda. This study involved the analysis of data from human subjects. Ethical approval for the CAfGEN protocol was received from the institutional and ethical review boards from all collaborating sites. Written informed consent and or assent were obtained from the participant’s authorized guardian/next of kin and all children from whom assent as required as per local ethical and regulatory guidelines appropriate for age.

### DNA extraction, targeted sequencing and allele identification for *KIR,* and HLA class I genotyping

DNA was extracted in Uganda and Botswana using standardized operating procedures and shipped on dry ice and liquid nitrogen to partner laboratories at the Baylor College of Medicine, Houston and the University of California San Francisco (UCSF), USA. We performed quality control and quantification of the DNA using the PicoGreen kit (Thermo Fisher Scientific, Waltham, MA); and determined the DNA fragment size using Nano drop and Bio-analyzer (Agilent, Santa Clara, CA). Samples were pooled and enriched for the *KIR* region following the previously described targeted capture protocol^10^. We performed targeted sequencing on the Illumina HiSeq 4000 (Illumina, San Diego, CA).

*KIR* alleles were determined using the updated version of our custom bioinformatics PING pipeline, which takes short-read sequence *fastq* files through multiple alignment, filtration and thresholding steps^37, 59^. *KIR* alleles were manually curated at a three-digit resolution to resolve the ambiguities. *KIR* allotypes were determined by searching the *KIR* Immuno-Polymorphism Database, IPD (https://www.ebi.ac.uk/ipd/kir/alleles) and the number of allotypes per gene was determined. KIR allotypes were defined as every unique protein coded for by a KIR allele. Potentially novel alleles were those identified in our data but had no *KIR* sequence found in the IPD-*KIR* database release 2.10.0^60^. HLA –A, –B, and –C alleles were classified according to their *KIR* interaction.

### Assessment of differential selection of *KIR* alleles

We compared the observed heterozygosity to the expected heterozygosity values for each allele obtained from GenAlEx software^61^. The Hardy Weinberg Equilibrium test was performed using the Guo and Thompson method in Python for population genetics analysis, PyPop software^62, 63^. The Ewens Watterson homozygosity test was performed for LTNPs and RPs in PyPop software^63^ using variances obtained by simulations of 10000 replicates to calculate the normalized deviate F_nd_ test^64^. This statistic denotes how much homozygosity is expected within a population under the Hardy-Weinberg equilibrium based on the exact test by Slatkin^65^. Statistical significance of the F_nd_ is denoted when the two-tailed P value is □<□0.025 or >0.975 using the Exact test^64^.

### *KIR*-HLA interaction and association analysis

We compared the frequencies of HLA-A, –B, and –C genotypes (at four-digit resolution) from our study with publicly available data at the same resolution from published a non-HIV North African cohort and a study from Sub-Saharan Africa^26, 29^. The presence of HLA allotypes per individual was assigned scores of 0 (none), 1, 2, 3, or 4 with each allotype counted as one irrespective of homozygosity. We performed an association analysis between KIR alleles and disease progression in PyHLA software considering a frequency threshold of 0.01, Fisher’s exact test, an additive model and multiple testing adjustments based on false discovery rate (FDR). A P value of less than or equal to 0.05 was considered statistically significant. The odds ratios and their 95 % confidence intervals were presented. All data visualizations were performed using R statistical software^66^.

## Results

### Participant description

We performed quality control for 349 samples in the PING pipeline with 6/260 Uganda and 22/89 Botswana samples dropped due to less than 50 reads being present, a minimum threshold for *KIR* allele calling. The 312 analyzed study participants were comparable by sex (in Uganda 52.1 % males vs 45.4 % males in Botswana). We also found that the Uganda cohort had significantly more LTNPs than RPs (148 Vs 25, P-value =0.001) as compared to the Botswana cohort (**Table 1**).

**Table 1:**
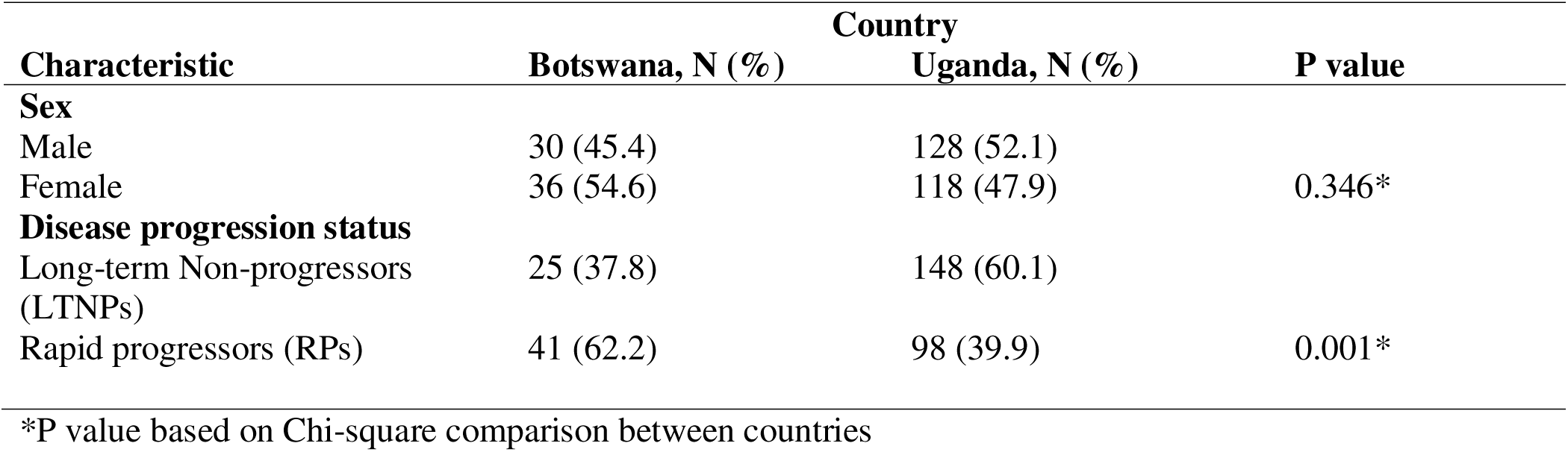
Demographic characteristics of the study participants.

### High allele diversity of the different *KIR* genes in our study cohorts compared to global populations

We evaluated the *KIR* allele diversity of participants in our study cohorts from Botswana and Uganda (combined sample, N=312) from allelic data generated using targeted sequencing and the updated PING pipeline. This allowed the characterization of the allele diversity to a three-digit level. The three-digit level resolution\ implies that the identified DNA sequence differentiates the allele by substitutions that change the protein sequence^67^. All 13 *KIR* genes were genotyped in our study cohort samples. We found that *KIR3DL3, KIR3DL1/S1*, *KIR2DL4* and *KIR3DL2* genes had many alleles identified in the entire cohort sample (**Figure 1 and Supplementary Table, S1)**. Notable were the few alleles identified for *KIR2DS1, KIR2DS2* and *KIR2DL5A* genes.

**Figure 1:**
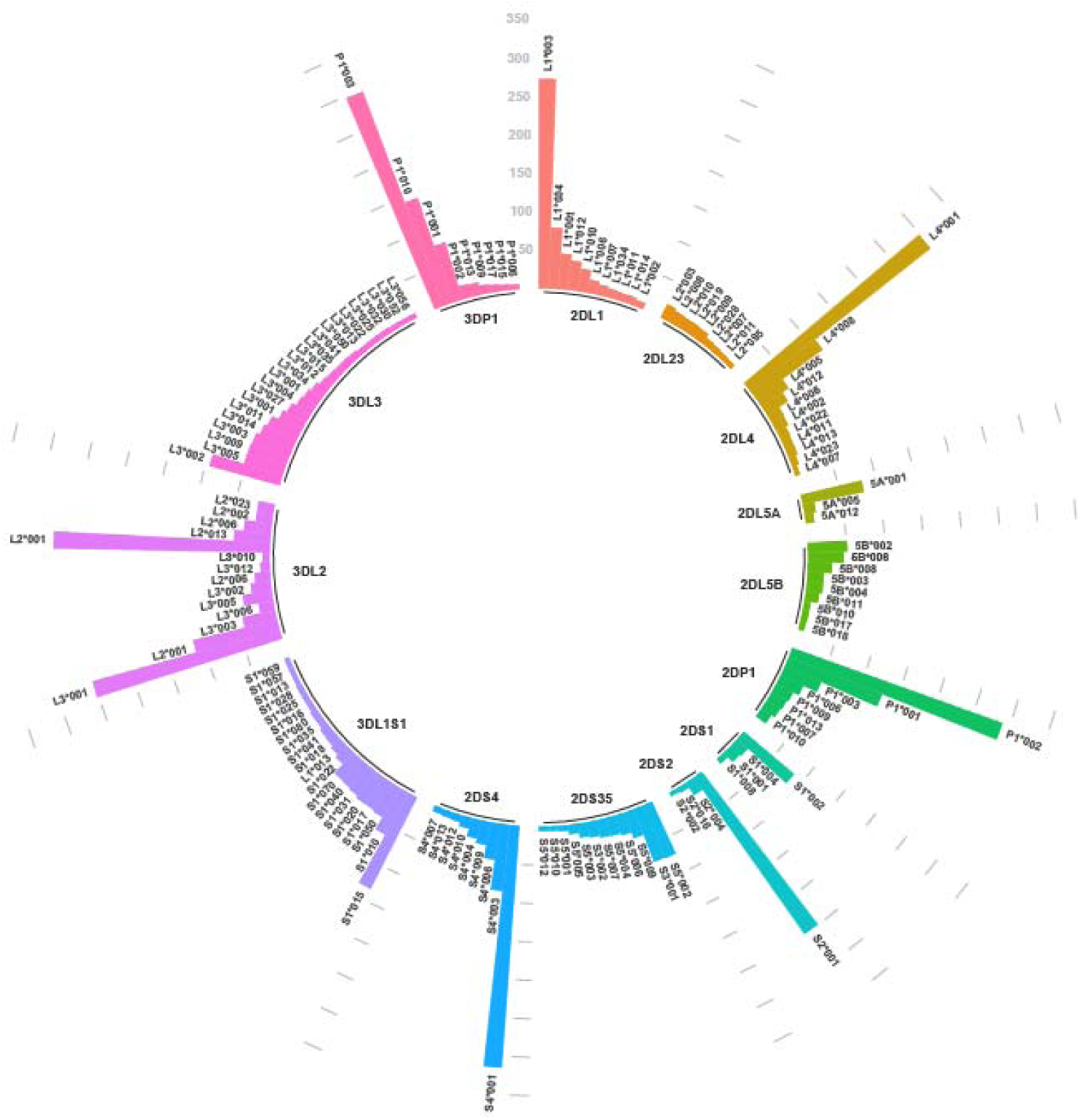
Circular bar plot of the diversity of the *KIR* alleles for the combined dataset of Uganda and Botswana (N=312). Only alleles with frequencies of more than 1 % are considered.

Based on the frequencies of the different alleles at the three-digit resolution, we compared our results with global populations that have evaluated *KIR* alleles. An allele frequency threshold of greater than or equal to 1% has been previously shown to be optimal for comparison across multiple studies with different sample sizes^24^. We found a high number of *KIR* alleles in the Ugandan population compared to the Botswana population (147 vs 111 alleles). On comparing the diversity of *KIR* alleles with global populations, a high number of alleles (average 129 ± 26) were found in our combined Botswana and Uganda cohort as compared to published data from global populations with approximately 73±13^24^.

Our study findings were comparable with sub-Saharan populations in terms of the average number of alleles (Namibian Nama: 100 vs Botswana& Uganda: 129; Tanzanian Hadza 66 vs Botswana and Uganda: 129) (**Supplementary Table, S2)**.

### High *KIR* allele diversity was observed in Uganda than in Botswana cohorts

We assessed for country-specific *KIR* allelic differences in 246 Ugandan and 66 Botswana samples, direct counting was used to determine the allele frequencies with the total number of alleles divided by 2N. In the Uganda cohort, *KIR2DS4*001* had the highest allele frequency of 0.502, followed by *KIR2DL1*003* (allele frequency of 0.496) and *KIR3DL2*001 (*allele frequency of 0.469). In the Botswana cohort, the *KIR2DL4*001* had the highest allele frequency (0.545) followed by *KIR2DS4*001* (0.515) and *KIR3DL2*001* (0.379) (**Supplementary Table, S3**).

We next determined whether the identified alleles were expressed and coded for any proteins (allotypes) identifiable in the IPD^60^. We also assessed if the identified allele was shared/found in both the Botswana and Uganda cohorts. The different inhibitory *KIR* genes exhibited a higher number of alleles than the activating *KIR* genes. We found that the Uganda cohort had more distinct alleles per gene as compared to the Botswana cohort samples. For example, for *KIR3DL2*, 40 alleles were identified in the Uganda samples as compared to eight alleles in the Botswana samples (**Table 2**). We also found that *KIR3DL3, KIR2DL1,* and *KIR3DL2* had 100% allotypes detected in both Botswana and Uganda. Overall, one of the framework genes, *KIR3DL3* had the highest number (n=12) of shared alleles between Botswana and Uganda.

**Table 2:**
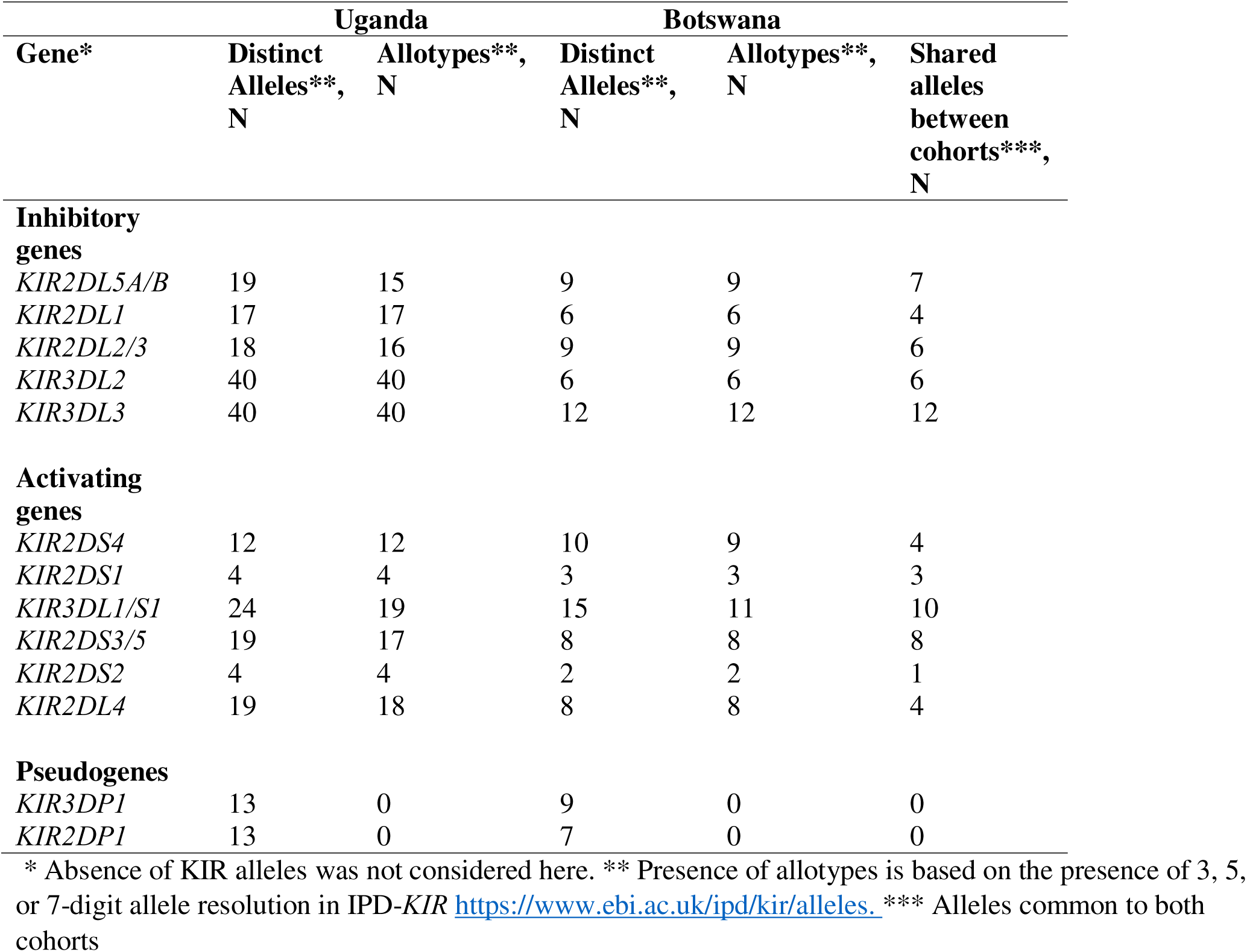
Number of unique alleles per gene identified in Uganda and Botswana cohorts.

| Gene* | Uganda |  | Botswana |  | Shared alleles between cohorts***, N |
| --- | --- | --- | --- | --- | --- |
|  | Distinct Alleles**, N | Allotypes**, N | Distinct Alleles**, N | Allotypes**, N |  |
| **Inhibitory genes** |  |  |  |  |  |
| *KIR2DL5A/B* | 19 | 15 | 9 | 9 | 7 |
| *KIR2DL1* | 17 | 17 | 6 | 6 | 4 |
| *KIR2DL2/3* | 18 | 16 | 9 | 9 | 6 |
| *KIR3DL2* | 40 | 40 | 6 | 6 | 6 |
| *KIR3DL3* | 40 | 40 | 12 | 12 | 12 |
| **Activating genes** |  |  |  |  |  |
| *KIR2DS4* | 12 | 12 | 10 | 9 | 4 |
| *KIR2DS1* | 4 | 4 | 3 | 3 | 3 |
| *KIR3DL1/S1* | 24 | 19 | 15 | 11 | 10 |
| *KIR2DS3/5* | 19 | 17 | 8 | 8 | 8 |
| *KIR2DS2* | 4 | 4 | 2 | 2 | 1 |
| *KIR2DL4* | 19 | 18 | 8 | 8 | 4 |
| **Pseudogenes** |  |  |  |  |  |
| *KIR3DP1* | 13 | 0 | 9 | 0 | 0 |
| *KIR2DP1* | 13 | 0 | 7 | 0 | 0 |

\* Absence of KIR alleles was not considered here. \*\* Presence of allotypes is based on the presence of 3, 5, or 7-digit allele resolution in IPD-KIR https://www.ebi.ac.uk/ipd/kir/alleles. \*\*\* Alleles common to both cohorts

We next explored the IPD-*KIR* to determine if there were potential novel alleles in our study cohorts. Novel alleles were defined as those that did not match any allele in the IPD-*KIR*. Six potential novel *KIR* alleles were identified in *KIR3DL1*/*S1* in the Ugandan cohort as compared to two novel alleles in the Botswana cohort (**Table 3**). *KIR2DL5A*/*B* has four potential novel alleles identified in Uganda as compared to none in Botswana.

**Table 3:** Potential candidate KIR genes in Uganda and Botswana cohorts.

| KIR gene | Potential candidate novel alleles* |  | Number of individuals |
| --- | --- | --- | --- |
|  | Uganda | Botswana |  |

|  |  |  | with possible<br>novel alleles** |
| --- | --- | --- | --- |
| *KIR2DL5A/B* | *KIR2DL5A\*002*<br>*KIR2DL5B\*001*<br>*KIR2DL5A\*010*<br>*KIR2DL5B\*012* | None | 2, 2,4, 11 |
| *KIR2DL1* | None | None | 0 |
| *KIR2DL2/3* | *KIR2DL3\*014* | None | 1 |
| *KIR3DL2* | None | None | 0 |
| *KIR3DL3* | None | None | 0 |
| *KIR2DS4* | None | None | 0- |
| *KIR2DS1* | None | None | 0 |
| *KIR3DL1/S1* | *KIR3DL1\*050,*<br>*KIR3DL1\*010,*<br>*KIR3DL1\*070,*<br>*KIR3DL1\*013,*<br>*KIR3DL1\*033,*<br>*KIR3DL1\*049* | *KIR3DL1\*010*<br>*KIR3DL1\*050* | Uganda -38,<br>39,26,16,3, 2;<br>Botswana-23, 6 |
| *KIR2DS3/5* | *KIR2DS35\*004*<br>*KIR2DS35\*009* | None | 1, 1 |
| *KIR2DS2* | None | None | 0 |
| *KIR2DL4* | *KIR2DL4\*101* | None | 0 |
| *KIR2DP1\*\*\** | - | - |  |
| *KIR3DP1\*\*\** | - | - |  |

\*Based on absence in the IPD KIR database. \*\*Based on the order of the alleles under the country columns, \*\*\*These are pseudogenes and do not code for any protein.

### The *KIR3DL2* gene shows significant positive selection in the Ugandan cohort

To explore the selection occurring with the *KIR* locus within the Uganda and Botswana cohorts, we first assessed the differences in the levels of heterozygosity in these study populations. The data show that Uganda had more observed heterozygosity than Botswana samples. The observed heterozygosity across the 13 *KIR* genes was less than expected overall. Considering a frequency threshold of 0.50, the observed heterozygosity in Uganda for *KIR2DL1, KIR2DS2, KIR2DL5, KIR2DP1 and KIR2DS3/5* was less than expected, indicating a possible positive directional selection process (**Figure 2A**). According to the data, the observed heterozygosity in the Botswana samples was zero for *KIR2DL1, KIR2DL2/3, KIR2DL4, KIR2DS1, KIR2DS2, KIR3DL2 and KIR2DP1* (**Figure 2B**).

**Figure 2A:**
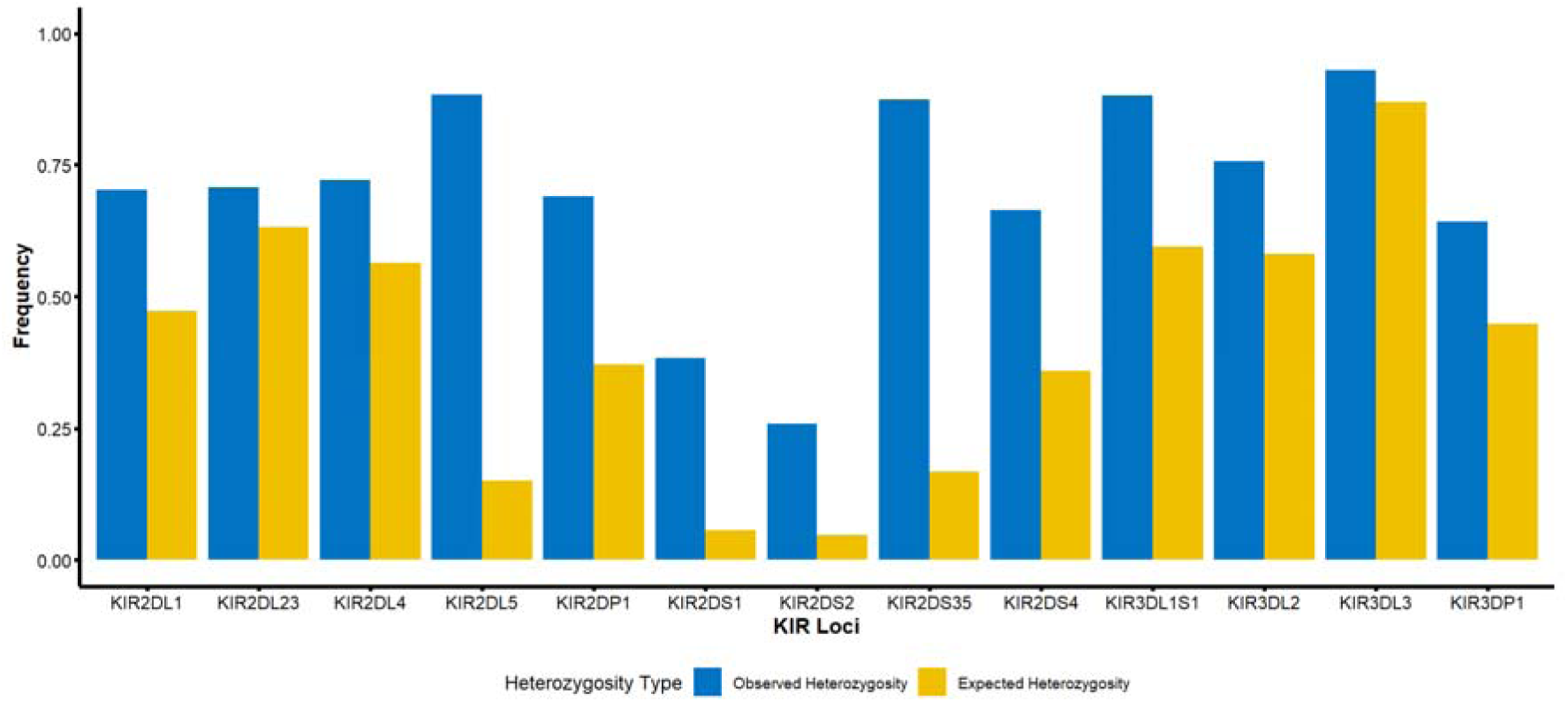
Overview of the comparison of the observed and expected heterozygosity across the 13 *KIR* genes in the Uganda cohort.

**Figure 2B:**
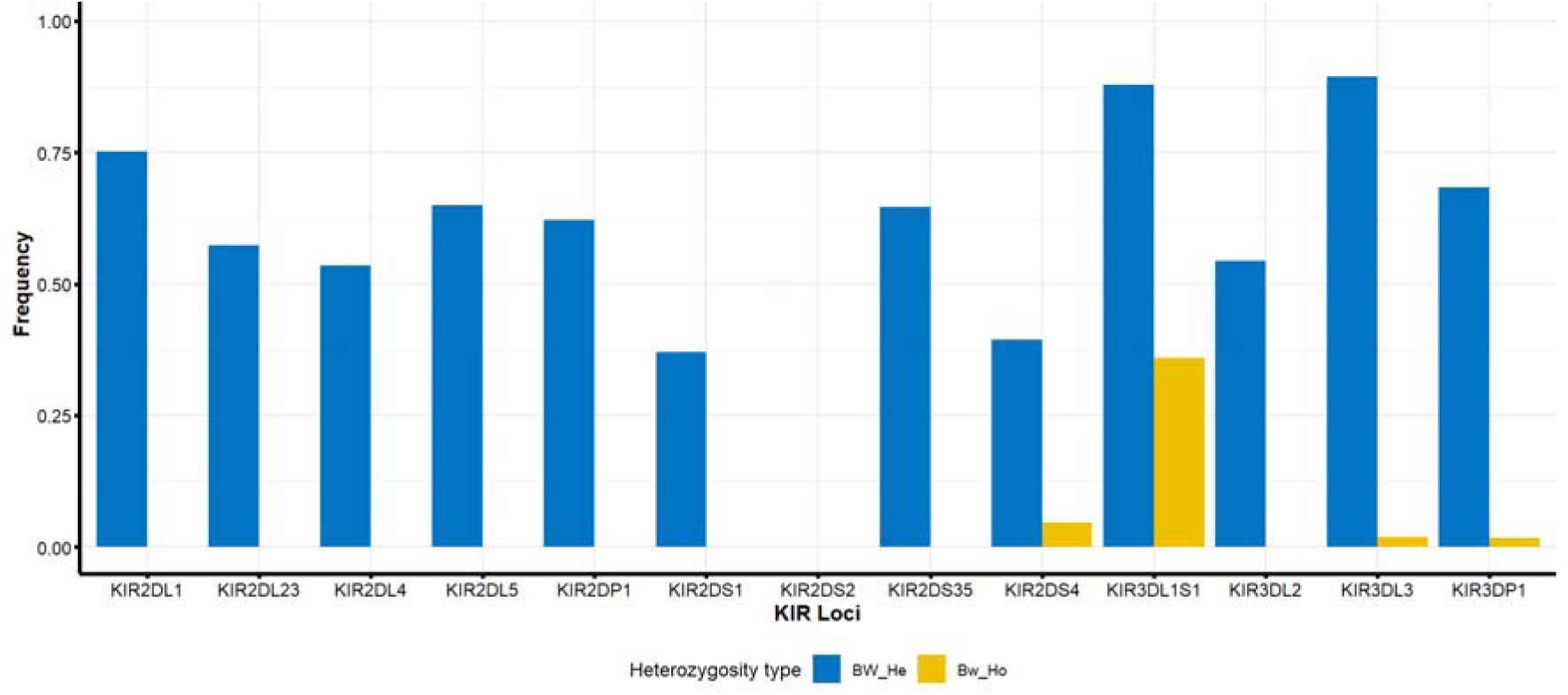
Overview of the comparison of the observed and expected heterozygosity across the 13 *KIR* genes in the Botswana cohort.

Since the Botswana samples had minimal diversity in this dataset (as evidenced by the observed heterozygosity) and lacked comparative MHC class 1 HLA data, subsequent analyses were performed considering **only** the Ugandan samples.

Next, we assessed the selection of the 13 *KIR* genes using the Ewens Watterson test in the Ugandan samples with consideration of the LTNP versus RP status. The Ewens Watterson uses a Monte-Carlo implementation of the exact test by Slatkin^65^. The reported P-values (against the alternative of balancing selection) are one-tailed or can be interpreted as two-tailed by considering the extremes (<0.025 or >0.975) of the null distribution of the homozygosity statistic under neutrality. In the Uganda cohort samples, there was a significantly high F_nd_ normalized deviate (F_nd_ positive values) for *KIR3DL2* among the study population, indicating a directional selection towards increased homozygosity of this region **(Figure 3).** The centromeric genes had more positive values than the telomeric genes except for telomeric *KIR3DL2* where LTNPs had slightly higher F_nd_ than RPs (LTNP F_nd_ = 4.2617, P-value = 0.9954 vs RP F_nd_ = 4.0273, P-value =0.9930). Although not statistically significant, there was marked heterozygosity of the *KIR3DL1/S1* gene with F_nd_ values higher among the LTNPs than the RPs ((LTNP F_nd_ = –0.5129, P-value =0.3416 vs RP F_nd_ = –0.4767, P-value =0.3686).

**Figure 3:**
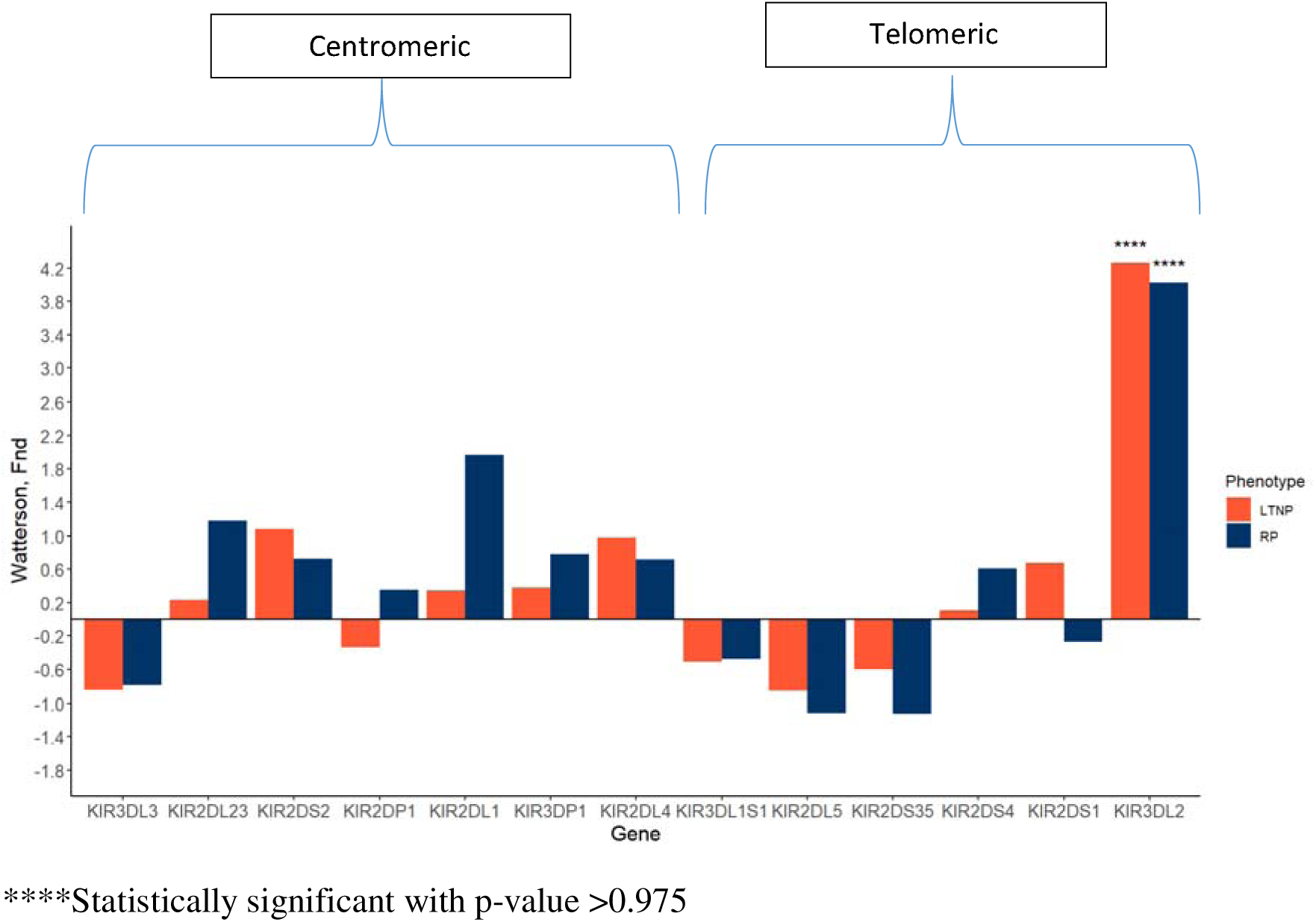
Level of homozygosity as estimated by Ewens-Watterson, F normalized deviate for Uganda samples.

### A high diversity of *KIR*-HLA ligand interactions was identified in the Uganda cohort

HLA and *KIR* segregate independently on different chromosomes and therefore an individual may have KIR but no ligand, and vice-versa^1^. This leads to functional and combinatorial diversity. We observed 61 HLA-A and HLA-B and 42 alleles of HLA-C in the Uganda samples (**Supplementary Table, S4)**. Regarding the phenotype status in Ugandan cohort, HLA C*03:02, C*07:01, C*06*02; HLA A*30:01, A*30:02; and HLA B*57:03, B*53:01, B*35:01, B*57:02, B*58:01, B*58:02 frequencies were higher in the LTNPs versus the RPs (**Supplementary Table, S4**) confirming and expanding our earlier findings^33^. We compared the HLA genotype frequencies in the Ugandan cohort dataset to a recently published non-HIV adult cohort in Egypt (both at four-digit typing)^29^, and we found that more HLA genotypes were found in the Ugandan sample than in the Egyptian data (**Supplementary Table, S5**). In comparison with another dataset that had Sub-Saharan non-HIV adult individuals^26^, there were similar frequencies of all identified alleles in both cohorts.

We were interested in *KIR*-HLA interactions because *KIRs* on NK cells work through their ligands to perform their effector functions^18^. HLA-KIR allotypes (based on amino acid sequence) were determined as described previously^68, 69^. HLA-A (A*23, A*24, A*25, and A*32) and HLA-B are highly Bw4-specific for *KIR3DL1* with some classified as 80I (Isoleucine) or 80T (Threonine) due to different amino acids present at position 80^52, 70^. The inhibitory *KIR2DL1* binds HLA-C group 2 allotypes with lysine at position 80, and *2DL2/3* and *KIR2DS1* receptors bind HLA-C group 1 allotypes, with asparagine at the same position^71^. *KIR2DS4* binds HLA-C1 and C2 alleles^72^ while *KIR3DL2* binds HLA-A*11/HLA-A*03^73^. Therefore, we assessed the differences in HLA allotype combinations within the Uganda sample cohort. By frequency, 52.3% of the HLA-C were C2 and 47.6% were C1. Among the HLA-B, 17 allotypes of *bw4*-80I were identified (**Supplementary Table, S4**).

Following the determination of HLA allotype combinations, the number of HLA allotypes carried per individual was examined. In the Uganda cohort, 103/246 (41.8%) of the individuals had two allotypes while 98/246 (39.8%) had three allotypes present (**Figure 4A**). On comparing the number of HLA-allotypes between our phenotypes, we found that LTNPs had higher frequencies of 2 and 3 allotypes when compared to RPs. (**Figure 4B)**. LTNPs had a higher frequency of HLA C1C2, C1C1 and C2C2 allotypes than RPs, although this was not statistically significant. However, the frequency of bw4-80I was higher among the LTNPs than RPs (8.9 % Vs 2.0%, P-value: 0.032) **(Table 4).**

**Figure 4A:**
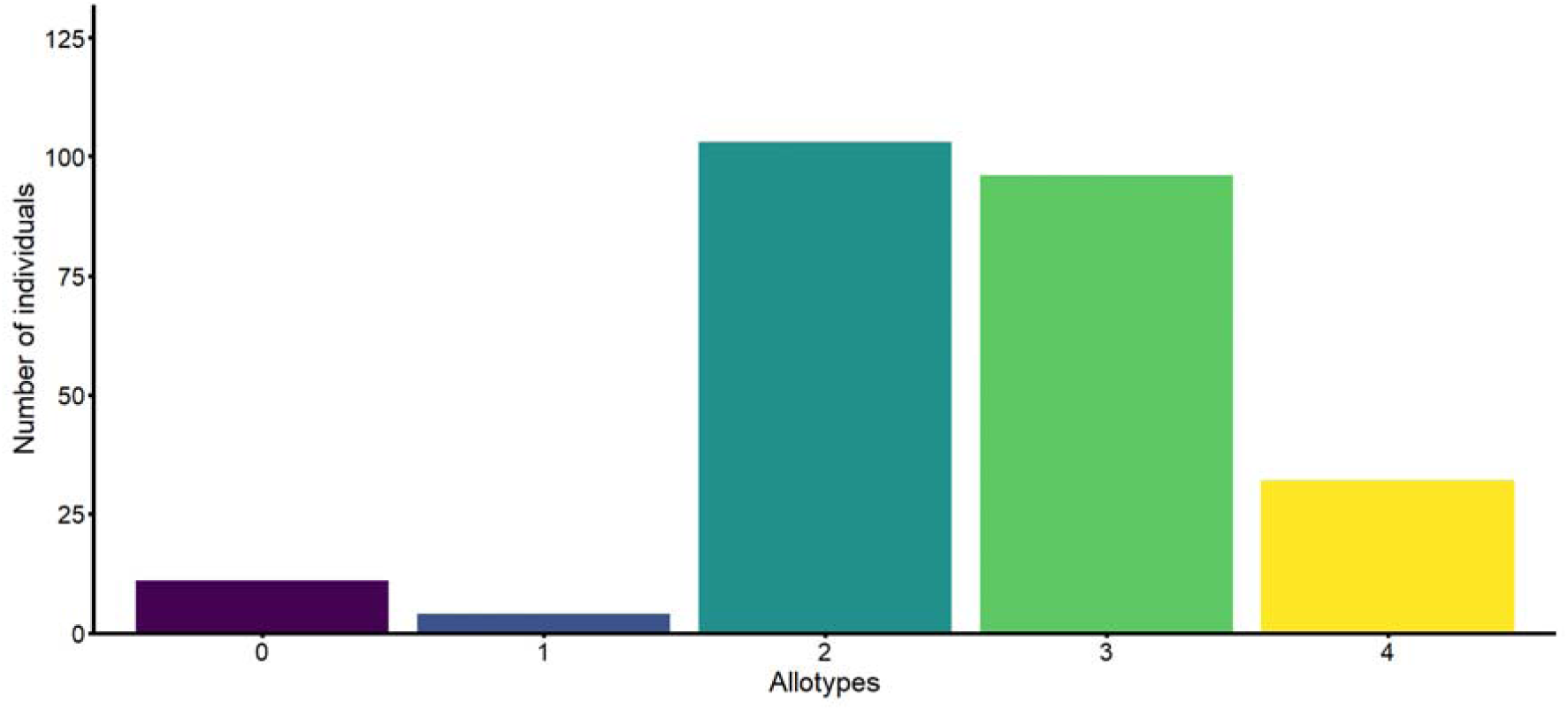
Bar plot showing the number of HLA allotypes per individual

**Figure 4B:**
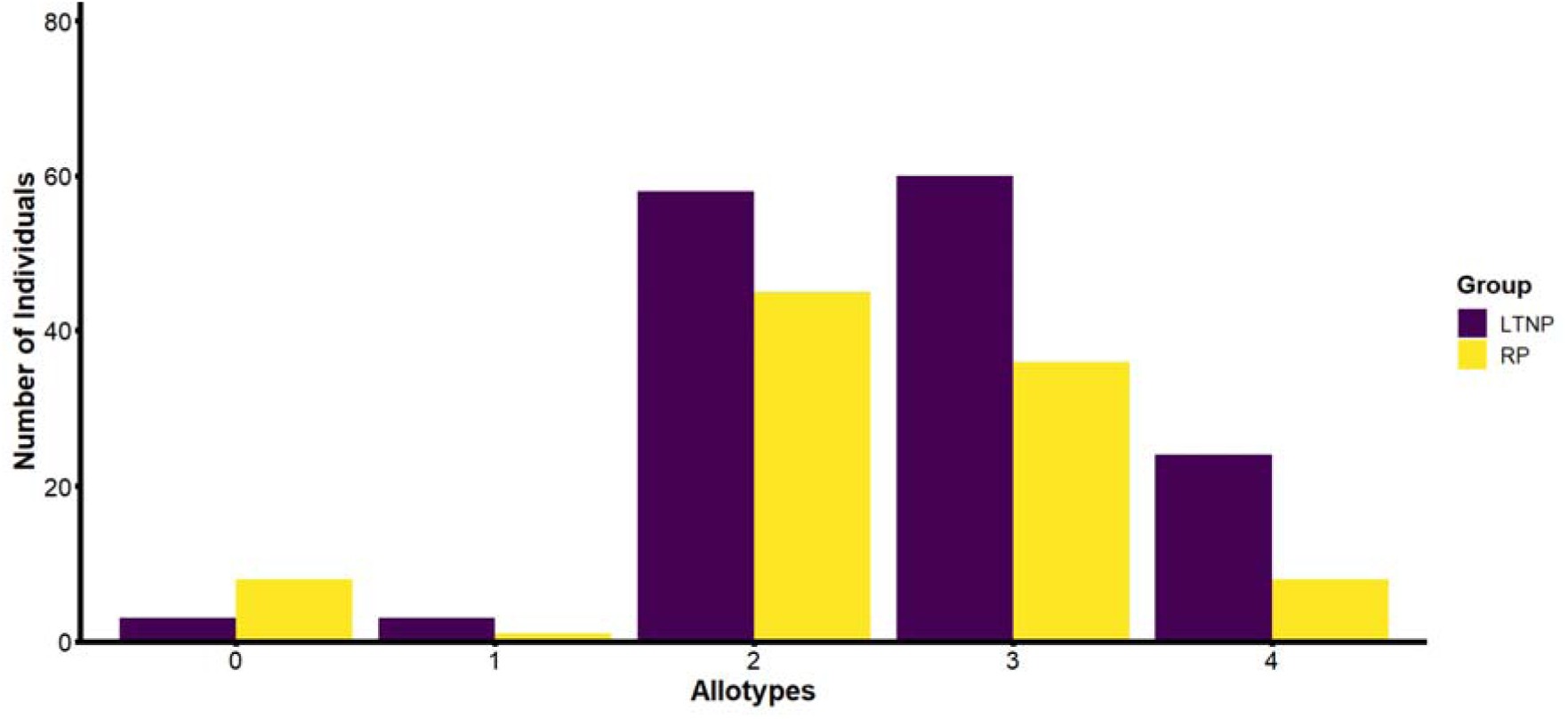
Bar plot showing the frequency of HLA allotypes per LTNP vs RP individuals in the Ugandan samples.

**Table 4:** Frequency of HLA-Allotypes between LTNPs and RPs.

| HLA Genotype | Rapid progressors, N (%)* | Long-term non-progressors, N (%)* | P values |
| --- | --- | --- | --- |
| C1C2 | 18 (7.8) | 31 (13.4) | 0.956 |
| C1C1 | 17 (7.5) | 26 (11.4) | 0.995 |
| C2C2 | 25(10.9) | 43(18.7) | 1 |
| Bw4-80I (HLA-Bw4) | 5 (2.0) | 22 (8.9) | 0.032 |
| A3/A11 | 13 (4.9) | 7 (2.8) | 0.054 |

\*column percentages

### *KIR2DS4*001* allele was potentially protective of long-term HIV disease non-progression

Previous research has documented the association between *KIR* allele and genotype frequencies and disease progression. For example, *KIR3DL1/S1* was associated with delayed disease progression^74^. For this analysis, we trimmed the Ugandan dataset from 246 to 203 individuals for whom no HLA genotype data was missing. We identified the *KIR2DS4*\**001* allele (OR: 0.671, 95 % CI: 0. 481-0.937, FDR adjusted P value=0.142) and *KIR2DS4*\**006* (OR: 2.519, 95 % CI: 1.085-5.851, FDR adjusted P value=0.142) were not significantly associated with slow HIV disease progression when considering an additive model under a logistic regression and the false discovery rate adjustment for multiple testing (**Table 5**). The rest of the model results are summarized in **Supplementary Table, S6**.

**Table 5:** Association between KIR alleles and HIV disease progression based on the additive model and logistic regression.

| Allele | Allele<br>LTNP | RP | Frequencies<br>Total | Unadjusted<br>P value* | Odds ratio (95 %<br>Confidence Interval) | P<sub>c</sub> |
| --- | --- | --- | --- | --- | --- | --- |
| ***KIR2DS4\*001*** | 0.435 | 0.5813 | 0.4926 | **0.019** | **0.671(0.481-0.937)** | **0.142** |
| *KIR2DS4\*010* | 0.0203 | 0.0312 | 0.0246 | 0.5273 | 0.688 (0.215-2.196) | 0.599 |
| ***KIR2DS4\*013*** | 0.0041 | 0.00437 | 0.0197 | 0.1003 | 0.228 (0.039-1.329) | 0.301 |
| *KIR2DS4\*003* | 0.2033 | 0.15 | 0.1823 | 0.2487 | 1.310 (0.828-2.074) | 0.448 |
| *KIR2DS4\*004* | 0.0285 | 0.0625 | 0.0419 | 0.1403 | 0.499 (0.199-1.256) | 0.316 |
| ***KIR2DS4\*006*** | 0.1098 | 0.0312 | 0.0788 | **0.0316** | **2.519(1.085-5.851)** | **0.142** |

\*Unadjusted P value based on logistic regression, P<sub>c</sub> –P value Based on false discovery rate

## Discussion

Previous research has documented the relevance of identifying the diversity of *KIRs* in understanding the HIV disease associations, population characterization and co-evolution with HLA ligands^25, 26, 29, 35^. The understanding of *KIRs* has also generated new insights into human genetic diversity on the African continent and their co-evolution with HLA ligands^26^. We robustly analysed the *KIR* diversity among populations from Uganda and Botswana using multiplexed next-generation sequencing techniques and new bioinformatics pipelines. To our knowledge, this is among the first extensive studies among children to identify and verify *KIR* allelic calls and HLA ligands. This study also extended our understanding of *KIR* and HLA allotypes among the Uganda cohort and elucidated the presence of positive selection for the *KIR3DL2* gene in this population. Our meticulous analysis of *KIR* genotypes revealed a significant proportion of HLA bw4-80I allotypes in RPs compared to LTNPs. We found that *KIR2DS4*001* was potentially protective against delayed disease progression.

Recombination complexities in the *KIR* genomic region lead to increased diversity of the *KIR* alleles and haplotypes. Analysis of the *KIR* allele frequencies in our study revealed that the centromeric region was more diverse in allele level content than the telomeric region. In comparison with other non-sub-Saharan African populations, increased diversity was observed in our study population^29^. Our study also shows that the *KIR* allele diversity extends to the differences between the Ugandan and Botswana cohorts with more diversity seen in the Uganda cohort. This aligns with published literature that the African region known as the cradle of humankind is highly heterogeneous in terms of ancestry, admixture and genetic diversity^30^ that extends this observation to the *KIR* region^25, 75^. We identified potential novel *KIR* alleles in the *KIR2DL5A/B* and *KIR3DL1/S1* genes in the Uganda and Botswana cohorts. These preliminary findings illustrate that in-depth analyses of *KIR*s have the potential to lead to discoveries of new alleles among African populations.

High *KIR* expression on the NK cell surfaces is associated with high NK cell reactivity^76^. We also found more alleles at higher frequencies in *KIR2DS4. KIR3DL2, KIR2DL4* and *KIR2DL1*. These findings point to possible high diversity within the region and reinforce previous findings concerning the *KIR* region^20, 29^.

The immune modulatory role of *KIR*s in infections like HIV and cancer has been underscored in previous research^42, 45^. Importantly, NK cells regulate antibody production to kill identified target infected cells^77^. We found more heterozygosity between the different *KIR* alleles in Uganda than in the Botswana cohort. From an evolutionary standpoint, high heterozygosity implies that the effects of genetic drift due to environmental exposures, intermarriages, and mutations within this region are still apparent and work in concert to lead to the fixation of alleles within the population^78^. Our findings of low heterozygosity in the Botswana cohort are contrary to findings of high diversity across the genome identified in a paediatric HIV cohort from Botswana^34^. The differences in findings may point to loci-specific variation and limitations in the breadth of *KIR*-specific Botswana reference sequences in the public databases that are utilized during computation.

We found that more genes deviated towards homozygosity in the centromeric *KIR* region than in the telomeric region. This finding is similar to results of selection in *KIR* genes amng the Amerindians and Ache populations^24^. The *KIR3DL2* gene showed strikingly high F_nd_ values with LTNPs having higher homozygosity than the RPs. Our further literature review found no significant roles of *KIR3DL2* in HIV pathogenesis. However, previous studies have found a possible role of *KIR3DL2* in maintaining high tumor burden by preventing the activation of cell death in adult T–cell leukemia and licensing pathogenic T-cell differentiation in spondylitis^79, 80^. There are several possible explanations for this result. The genes in the telomeric region could be undergoing purifying selection as compared to the centromeric genes. HIV itself could also exert selection pressure differentially on the telomeric than centromeric region^81^. Similarly, given that the study cohorts were drawn from settings with high infectious disease burdens, we cannot rule out the other possible roles of infections like malaria and hepatitis in selection in our study. In contrast to earlier findings among the Amerindians, however, no significant evidence of selection among the *KIR2D2/3, KIR2DS35, KIR2DL5B* and *KIR2DL1* genes was detected in our study cohorts^24^. Among the Japanese, telomeric KIR genes were found to have balancing selection unlike our study findings^18^. Probable causes of the differences in findings may include genetic ancestry and exposure to different pathogens.

To perform the effector functions, *KIR*s utilize the HLA class I ligand molecules and have co-evolved across populations^26, 70, 72^. In our study, we found a high predominance of HLA-C2 allotypes as compared to HLA-C1 allotypes (c1=43.7%) in our Ugandan cohort. There are similarities in findings between the present study and those described by Nakimuli *et al* (C2=54.9 % and C1= 45.1 %) among healthy female Ugandan donors^35^. Other sub-Saharan African populations have demonstrated similar trends of higher HLA-C2 frequencies^82, 83^ unlike North African^29^ and non-African populations^24, 84^. Of note, we found significant bw4 allotype diversity differences between LTNPs and RPs in the Ugandan cohort. These findings though preliminary need further exploration as bw4-80I is a key ligand for the activating *KIR3DL1/S1* receptor that affects many disease phenotypes.

An emerging issue from our study was the potential protective association between *KIR2DS4*001* and long-term HIV disease progression where individuals with this allele were at lower odds of being LTNPs. This finding is a paradox given that the known activating effects of this *KIR2DS4* allele on the NK cells may mediate activated cell destruction thereby increasing the chances of an individual being an LTNP. *KIR2DS4*001* has also been associated with HIV transmission among sero-discordant couples in Zambia and altered HIV-1 pathogenesis among HIV-positive American youth in previous adult studies^85, 86^. However, there remains limited literature regarding *KIR2DS4* allele diversity and HIV phenotypes among the pediatric HIV populations. Intriguingly, no *KIR2DL2/3* allele was associated with HIV disease progression in our cohort although it is postulated that it segregates together with *KIR2DS4* on the telomeric haplotypes due to linkage disequilibrium. *KIR2DS4*006* has not been associated with any HIV phenotypes and its immunologic role remains unknown despite individuals with this allele in our study having higher odds of being LTNPs^87^. We did not find any statistically significant associations between *KIR3DL1/S1* and disease progression in our study; a deviation from previous studies^40, 88^. A limitation of determining phenotype associations in smaller sample sizes is the risk of random error and limited variability in the samples to detect differences. For this reason, we were not able to identify significant associations between other *KIR* alleles and HIV disease progression after correcting for multiple comparisons. We propose that *KIR* associations and mechanistic studies should be further explored in larger sample size and haplotype block analyses given the high number of genes and alleles at the *KIR* loci that are compared at once.

Our study is among the few studies that have evaluated the diversity of *KIR* alleles on the African continent among large HIV populations and expands evidence about the complexity of the region at the three-digit allele level. In the era of increased bioinformatics and availability of NGS techniques, our findings advance knowledge regarding the diversity of *KIR*, HLA alleles and their allotypes among African paediatric population samples and reduce the global disparities in *KIR* genomic research among diverse populations.

These data must be interpreted with caution because the study had some limitations. We found a high number of ambiguous allele calls in our samples. The ambiguity may arise due to multiple segments of unusual similarity created by recombination events in the *KIR* region, high refinement of the *KIR* genes database and nomenclature, unmatched primers for regions and base pair combinations in genes and phasing limitations due to the lack of parental sequence data^15^. Discrepancies may also arise due to few numbers from the sample population in the reference panels to decipher KIR alleles. However, we minimized this with resolution using manual techniques based on known standards of resolution of ambiguity. We did not perform follow-up Sanger sequencing to confirm the novel *KIR* alleles at the nucleotide level. We had a limited sample size for the association analyses between *KIR* alleles and HIV disease progression, which may have led to many insignificant results after correction for multiple tests. However, our results lay the groundwork for the discovery of a possible association between *KIR* and HIV in paediatric and adult African population settings. The Ewens Watterson test has limitations in that it determines recent selection as compared to other tests like the comparison of differences between non-synonymous and synonymous substitutions^89^ which evaluate selection at longer time scales. Our *KIR* and HLA diversity analyses did not have a non-HIV infected population from the same populations to decipher if the findings were specific to HIV. We also did not have an adult HIV+ comparison group from the same population, and we do not know if age (or the difference between perinatal and adult-acquired HIV) influences the findings.

In conclusion, we have presented an in-depth study of *KIR* diversity, *KIR* and HLA class I ligands, and associations with HIV disease progression among pediatric populations from Uganda and Botswana. Our findings point to a high diversity of *KIR* alleles among our Ugandan and Botswana cohorts as compared to global populations. Our study further demonstrates a positive directional selection of the *KIR3DL2* gene among Ugandan samples.

*KIR2DS4*001* and *KIR2DS4*006* were potentially associated with HIV disease progression. We recommend further research in the replication of our findings and functional validation of the *KIR* and HLA alleles’ associations with disease phenotypes.

## Supporting information

S1-The KIR allele frequencies in Uganda Botswana (overall), N=312), S2Comparison of alleles with global populations, S3-Frequency of the KIR alleles

## Acknowledgements

We would like to acknowledge the following members who are part of the CAfGEN consortium: Bhekumusa Lukhele, Edward D. Pettitt, and Marape Marape, who were participating co-investigators. We acknowledge Bathusi Mathuba, Nasinghe Emmanuel, Eddie Wampande, Thembela Mavuso, Buhle Dlamini, Abhilash Sathyamoorthi, Yves Mafulu, Harriet Nakayiza, Bheki Ntshangase, Keboletse Mokete, Kennedy Sichone, Keofentse Mathuba, LeToya Balebetse, Muambi Muyaya, Nancy Zwane, Nicholas Muriithi, Sibongile Mumanga, Thobile Jele, Olekantse Molatlhegi and Thato Regonamanye. We also acknowledge the staff of the Hollenbach lab, at UCSF, USA, the Childhood Complex Disease Genomics lab at the National Human Genome Research Institute, USA and the African Center of Excellence in Bioinformatics and Data Intensive Sciences at the Infectious Diseases Institute, Kampala, Uganda for offering computational support and guidance. Finally, we thank the children and their caregivers in Uganda and Botswana who participated in the study.

## Funding

The grant under award Number, U54AI110398 administered by the National Institute of Allergy, supported the project described and Infectious Disease (NIAID), Eunice Kennedy Shriver National Institute of Child Health & Human Development (NICHD), and National Human Genome Research Institute (NHGRI) as part of the NIH Common Fund H3Africa Initiative. Additional funding was obtained from the Nurturing Genomics and Bioinformatics Research Capacity in Africa (BRECA) grant number # U2RTW 010672 from the NIH Fogarty International Center. The funders had no role in the design, interpretation and publication of the study findings

## Conflict of interest

The authors declare no conflict of interest.

## Author contributions

JM, SK, MA, EK, GM, TD, GS, SM, GM, GR, LW, BM, MM, DJ, DPK, MLJ, NH and JAH, conceptualization, writing – review and editing, funding acquisition, investigation, and project administration. JM, SK, MA, SM, NH, and JAH: data curation. JM, SK, DJ, and JAH: formal analysis. MLJ, GM and MM: funding acquisition. and JK-L: investigation. JM, GM, MA, GS, SM, GM, NH, and JAH: methodology. DJ, GM, NH, MLJ, MM: project administration, GM, NH and JAH: supervision. JM, SK, NH and JAH: validation and writing – original draft. All authors contributed to the article and approved the submitted version.

## Data availability

Datasets used in this paper are available in online repositories with accession numbers: https://h3africa.org/wp-content/uploads/2018/05/App-D-H3Africa-Data-and-Biospecimen-Access-Committee-Guidelines-final-10-July-2017.pdf, NA.x

## Abbreviations

FDR: False Discovery rate
*KIR*: Killer cell immunoglobulin-like receptor
IPD: Immuno-Polymorphism Database
HIV: Human Immune Deficiency Virus

## Supplementary information

Supplementary tables are provided in the online version of this manuscript

| Item | Description |
| --- | --- |
| **Supplementary Table S1** | The KIR allele frequencies in Uganda Botswana (overall), N=312 |
| **Supplementary Table S2** | Comparison of alleles with global populations |
| **Supplementary Table S3** | Frequency of the *KIR* alleles in the Ugandan (Upper) and Botswana Cohorts compared by the Long-term and Rapid progressors phenotypes |
| **Supplementary Table S4** | Overall and comparative (LTNP versus RP) frequencies of the HLA alleles at four-digit resolution in the Ugandan Cohort |
| **Supplementary Table S5** | HLA class I allele frequencies from the Ugandan cohort (2N=496) compared with data from Sub-Saharan Africa (2N=366) and Egypt (2N=250) |
| **Supplementary Table S6** | Results of logistic regression with an additive model and allele frequency threshold of 0.01. |

## Notes

### Competing Interest Statement

The authors have declared no competing interest.

